# Computational fluid dynamics enables predictable scale-up of perfusion bioreactors for microvessel production

**DOI:** 10.64898/2026.03.24.713992

**Authors:** Pouyan Vatani, Kasinan Suthiwanich, Zidong Han, David A. Romero, Sara S. Nunes, Cristina H. Amon

**Author notes:** P.V., D.R. and C.A. developed the CFD models and conducted all in silico scale-up studies, using porosity and permeability values estimated by Z.H. based on experimental data. K.S. and S.N. performed all the microvessel network formation experiments on both platforms. All authors participated in drafting the manuscript, while D.R., S.N. and C.A. reviewed and approved the final manuscript.

## Abstract

Scaling up microvessel culture systems is essential for producing vascularized clinically relevant tissues, yet current platforms offer little guidance on how to preserve flow conditions during scale-up. Here, we present a computational–experimental framework using computational fluid dynamics (CFD) to guide the design and scaling of microvessel bioreactors. Interstitial flow distributions were predicted in two perfusion-based platforms–a permeable insert and a rhomboidal microfluidic chamber–across multiple scaling factors and hydrostatic pressures. CFD identified IF ranges conducive to vasculogenesis and quantified how geometry and pressure modulate flow uniformity. Scaled-up bioreactors generated microvessel networks with consistent morphology and connectivity over a 30-fold increase in culture volume, confirming that maintaining equivalent IF ensures reproducible outcomes. The permeable insert platform maintained uniform IF across scales, while the rhomboidal chamber produced spatially varying IF resulting in heterogeneous but physiologically relevant networks. These findings establish CFD as a predictive tool for rationally scaling perfusion bioreactors, enabling microvessel production at clinically relevant scales with controllable morphology.

**Significance Statement:** Scaling up microvessel bioreactors is critical for engineering large pre-vascularized tissues. However, larger scales may disrupt flow conditions that drive vessel formation. This study demonstrates that computational fluid dynamics (CFD) can predict interstitial flow and guide the rational scale-up while preserving the vasculogenic microenvironment. Experiments across 30+-fold size increase confirmed that matching inter-stitial flow results in morphologically identical microvessel networks. By linking simulation-based design with experimental validation, this work establishes CFD as design tool for scalable perfusion bioreactors for production of microvessel networks at clinically relevant scales.

## 1. Introduction

The vascular system enables the transport of oxygen and nutrients and the removal of waste products—functions essential for sustaining any living tissue (1). In engineered constructs, vascularization is critical for recapitulating these functions, as oxygen and nutrient diffusion into tissues is limited to approximately 200 µm; beyond this range, cells encounter hypoxia, lack of nutrients, and consequently undergo necrosis (2, 3). When an engineered construct is transplanted into a body, the rate at which the host vascularizes the graft can be as low as tens of microns per day (e.g., 120 µm*/*d (4)), resulting in inadequate oxygen and nutrient supply to the graft. To overcome this limitation, the preferred strategy is to integrate vasculature into the tissue grafts prior to its application, a process referred to as pre-vascularization (3, 5).

Engineered tissues are commonly pre-vascularized using co-culture of the target tissue cells with endothelial cells (ECs) derived from pluripotent stem cells (PSCs) or isolated directly from tissue sources (6). Co-cultures also include supporting cells, such as mesenchymal stem cells (MSCs), fibroblasts, or pericytes, which promote vascular stabilization and maturation by providing structural support and signaling to endothelial networks (6, 7). While co-culture methods have demonstrated success, recent studies show that incorporating pre-formed microvessels–such as those in micro-sized aggregates or microspheres (3, 5)–can accelerate perfusion and vascular integration upon implantation. These prevascularized constructs facilitate early perfusion by forming anastomoses with the host vascular system, a process that is 2X-3X faster than neovascularization (8–10).

During *de novo* formation of vascular networks *in vivo*, the microphysiological environment exposes endothelial cells to various biochemical and mechanical stimuli, such as shear stress, Extracellular Matrix (ECM) stiffness, pressure gradients, and growth factors (11, 12). In order to replicate these conditions, micro-physiological systems (MPS) have been utilized to study and modify engineered microvascular networks (13, 14) and micro-lymphatic networks (15). MPS have shown significant potential in tissue engineering applications, due to their capability to closely recapitulate *in vitro* the bio-chemomechanical environment(16) at the length-scales relevant for the target tissues. For vasculogenesis applications, MPS must integrate media perfusion, enhancing nutrient and waste transport while introducing controllable flow-induced mechanical stimuli, which has been shown to be key for the alignment and organization of endothelial cells into functional, perfusable microvascular networks (13, 17, 18). Platforms for vessel-on-chip applications have been designed in various configurations, such as idenTx, IFlowPlate, and customized open microfluidic devices (19–21).

A common design choice for vessel-on-chip applications is the use of one or several chambers containing cell-seeded fibrin or collagen gels, connected to two microfluidic channels running in parallel along the outline of interconnected chambers. The channels, operating at different pressures, induced flow through the chambers to provide nutrients, growth factors, remove waste, and exert the required shear stress. The chambers, usually designed with corrugation such as multiple-diamond or saw-teeth layouts, provide effective control and confinement of cell-embedded hydrogel, but result in a non-uniform interstitial flow (IF) field with both streamwise and cross-streamwise velocity components, and with larger velocities near the midline of the chambers. For instance, Jeon et al. (22) and Zervantonakis et al. (23) used such platforms to study angiogenesis, intra- and extravasation phenomena during in vitro tumour formation and metastasis, reaching interstitial flow velocities of 220 µm*/*s and shear stress of 0.025 Pa (22). A conceptually similar version of this platform featuring three interconnected rhomboidal chambers was used by Wang et al. (24), who reported microvessel formation throughout the chamber, with vessel diameters of approximately 50 µm near the chamber inlet and outlet. In a related study using the same rhomboidal chamber geometry, Hsu et al. (25) achieved a total vessel length of approximately 16 mm after two weeks of culture.

An alternative device design used to impose IF effects on endothelial cells uses permeable inserts on standard well plates, which facilitates vessel retrieval and offers design simplicity. Deng et al. (12) demonstrated the effectiveness of this platform by placing 12 mm column of growth media placed on top of a transwell with endothelial-seeded fibrin gel, resulting in microvessels with diameters ranging from 10.0 µm to 40.0 µm.

Current MPS systems for microvessel production–such as those described above–have demonstrated high physiological relevance in research settings. Notwithstanding these successes, they have a limited yield due to their small sizes, with tissue chambers of 0.2 mm^3^ and 25 mm^3^ for the rhomboid chamber and Permeable Insert (transwell) devices, respectively, and their scalability to support larger tissue constructs remains unproven. As tissue engineering applications move from the bench to the clinic, yield requirements for target cell types increase significantly (26). It has been shown that engineered heart tissues can be generated using a combination of 70% cardiomyocytes, 15% cardiac fibroblasts, and 15% endothelial cells, requiring as few as 1.6 ×10^4^ cells for small constructs(27). Scaling these ratios to clinically relevant sizes involves millions of cells and larger bioreactor volumes. For instance, tubular engineered heart tissues have been successfully fabricated with an inner diameter of 6 mm, wall thickness of 1 mm, and length of 18 mm, using cardiomyocytes at a density of 18.0 ×10^6^ cells/mL (28), requiring approximately 18.0 ×10^6^ endothelial cells/mL. Generating microvessels from these endothelial cells would require sustaining target IF conditions in up to 77 permeable insert platforms.

It is thus paramount to gain a thorough understanding of the scalability of these platform designs, the variations in flow-induced mechanical environment between different scaling strategies, and their effect on the yield and functional characteristics of resulting microvessel networks. Computational fluid dynamics (CFD) simulations enable this understanding by predicting the IF field for any device and chamber shape or size. When combined with in vitro microvessel formation experiments, CFD can enable richer characterization, interpretation and cost-effective optimization of perfusion-based platforms. For instance, Wang et al. (29) demonstrated through CFD simulations that a microfluidic platform design ensured uniform interstitial flow (pressure drop of 5 mm H_2_O, or 49 Pa) conducive to vasculogenesis and predictable cell flow paths. Similarly, Hsu et al. (25) used CFD to validate the ability of their platform to replicate physiological mass transport conditions with precise control over interstitial flows (Péclet numbers ranging from 0.0056 to 160). Hajal et al. (30) further explored the influence of flow conditions, both measuring (*in vitro*) and predicting (*in silico*) interstitial velocities between 0.08 µm*/*s and 2.14 µm*/*s, with predicted shear stresses of 0.15 Pa to 1.0 Pa, while elucidating the effect of luminal, transendothelial and interstitial flow conditions on tumor cell extravasation and migration. In summary, CFD models validated with *in vitro* data–such as those described above–can provide valuable insights for analysis and design of bioreactor platforms that create environments consistent with proliferation, differentiation, and maturation of endothelial cells and microvessel formation.

In this paper, we study the scalability of interstitial flow (IF) conditions across two distinct platform designs: Rhomboidal chambers and permeable insert setups. We investigate the feasibility of scaling up these designs while maintaining interstitial flow velocities within the physiological range of 0.1 µm*/*s to 11.0 µm*/*s (31), subject to constraints on the maximum height of a fluid column driving the perfusion flow. This size constraint effectively limits the maximum inlet pressure. To this end, we use CFD methods to predict, analyze and compare the IF fields within the culture chambers. Experiments conducted with these two platforms provide evidence regarding the ranges of IF that lead to successful formation of microvessel networks. In addition, comparative experiments with different platforms but similar average IF show that regardless of chamber shape or size, the characteristics of resulting microvessel networks are primarily dictated by the detailed IF field within the fibrin gel scaffold.

## 2. Materials and Methods

### A. Platform Designs

Two platforms were analyzed in this work, a *permeable insert platform* (12) and a *rhomboidal chamber platform* (29), which will hereafter be referred to as Platform A and Platform B, respectively, shown in Fig. 1. For simulation purposes, computer-aided design models (CAD) of the platforms were created in *SolidWorks* and subsequently imported into the *COMSOL* simulation software.

**Fig. 1.**
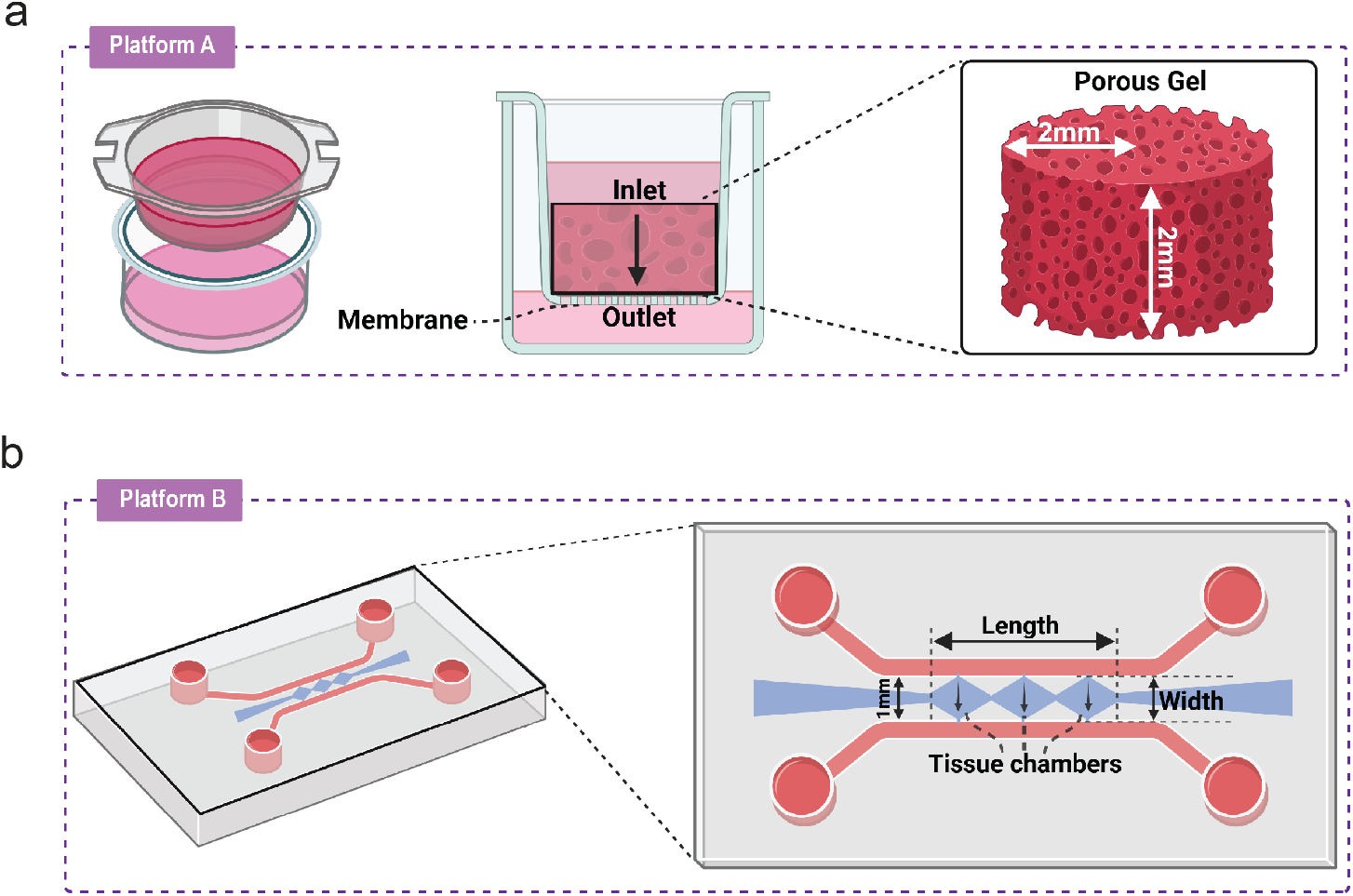
Microvessel network benchtop platforms. (a) Permeable insert (Platform A), including a 3D view (left), cross-sectional view (center) and corresponding computational simulation domain (right). A fluid column is placed on top of the inlet, exerting a pressure proportional to the column height. Media flows through the porous gel, through the porous membrane (8 μm pores) in the bottom and the outlet into the outer well. (b) Rhomboid chambers (Platform B), including a 3D view (left) and top view (right).

Scaled versions of the platforms were created by modifying the computer-aided design (CAD) models to allow key dimensions to be scaled independently or in combination. For Platform A, the gel chamber thickness (distance between inlet and outlet) and the chamber radius were the primary scaling parameters. For Platform B, the length (across all three rhomboidal tissue chambers) and the width (distance between inlet and outlet channels) were used. For any of the platforms, geometrical parameters were first scaled by factors of 2, 5, 10, and 20, applied independently to each dimension. Secondly, the same scaling factors were applied simultaneously to all dimensions scaling the entire geometry while maintaining the aspect ratio.

### B. Mathematical Model and CFD Simulations

The simulation models used in this work assume that the culture gel behaves as an isotropic, spatially homogenous porous media (29). Based on the pore dimension of the fibrin gel (32) and the average flow velocities through the gel under the conditions tested (approx. 5 µm*/*s to 20 µm*/*s) (29), the Reynolds number for this flow is (Re ≈ 8.45 ×10^−6^ to 3.38 ×10^−5^) (Re ≪1). Hence, we model the flow through the gel using Darcy’s Law for flow in porous media (33). The governing equation for interstitial flow is thus expressed as:

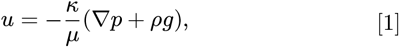

where *u* is the fluid velocity [m/s], *κ* is the permeability [m^2^], *µ* is the fluid viscosity [Pa·s], ∇*p* is the pressure gradient [Pa/m], and *ρg* is the gravitational force per unit volume [N/m^3^].

Darcy’s Law was solved together with the differential form of the Conservation of Mass principle, which for an incompressible porous media is:

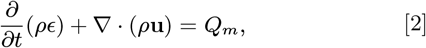

where *ϵ* is the porosity, and *Q_m_* accounts for any mass source or sink (assumed zero in this work). The solution of these equations provides interstitial flow velocity profiles within the gel, defined as the total volumetric flow rate through the bulk gel divided by the cross-sectional area of the gel. Thus, IF is a measure of an average, equivalent flow speed through a conduit of the same cross-sectional area but without the porous medium. Alternatively, the flow field inside the porous media can be expressed in terms of the average pore velocity, defined as:

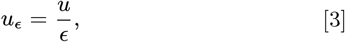

which represents the average flow velocity that any cells located inside the porous media pores will be directly subjected to. In this work, results are expressed in terms of interstitial flow (IF) to remain consistent with prior studies (34). However, the volume-averaged pore velocity (*u_ϵ_*) may offer a more physically relevant quantity for defining the flow conditions that are favorable to microvessel formation.

Equations 1 to 3 indicate that permeability and porosity are two critical properties of the porous media that modulate interstitial flow. Based on previous experimental data (35) and image analysis on dextran flow experiments (described in the supplementary materials), permeability and porosity values of 1.5 ×10^−13^ m^2^ and 0.91 were used in this work to generate all results.

The equations were discretized using the finite element method in COMSOL Multiphysics. The simulations were performed in a 2D axisymmetric model for Platform A and a 2D model for Platform B. A stationary solver with an iterative method was used. The convergence threshold was set to a mass conservation residual of 10^−15^, starting from initial residuals in the range of 10^4^. The computational domain for both platforms was meshed with triangular elements, and a mesh independence study was conducted to ensure that the results were not influenced by further mesh refinement. No-slip boundary conditions were enforced at chamber walls on both platforms. For Platform A, the pressure gradient was defined across the thickness of the gel, while for Platform B, the pressure drop was applied across each rhomboid-shaped tissue compartment, i.e., between the two microchannels.

### C. Microvessel network generation

Immortalized GFP-expressing human umbilical vein endothelial cells (HUVEC, passage 15–25, a gift from Dr. Roger Kamm, MIT (36)) were cultured in a completed VascuLife medium (LifeLine Technology) with only 25 % of the provided supplement volume of heparan sulfate. Normal human lung fibroblasts (NHLF, passage 5–6, Lonza) were cultured in DMEM (Gibco) supplemented with 20 % fetal bovine serum (heat inactivated, Wisent) and 1 % penicillin/streptomycin (Gibco). Human Brain Vascular Pericytes (HBVP, passage 4–6, ScienCell) were cultured in a completed Pericyte Medium (ScienCell).

The generation of microvascular network in Platform A was based on a previously established protocol (12). Briefly, cell-culture inserts (8 µm pore size) were modified with either a 2 mm-high PDMS ring (for 24-well sized insert) or PDMS film coating on the insert inner wall (for 6-well sized insert) to limit the hydrogel height and meniscus. All inserts were coated with poly-D-lysine (Gibco) to ensure hydrogel adherence to the insert. Dissociated HUVEC and stromal supporting cells (either NHLF or HBVP) were resuspended in a fibrinogen solution at 8–10 million cells/mL and 0.25–2 million cells/mL respectively, depending on specific experiments. With a pre-chilled pipette tip, cell-embedded fibrinogen solution was quickly mixed with thrombin in DPBS until the final mixture contained 10 mg*/*mL fibrinogen and 10 U*/*mL thrombin. The resulting hydrogel was immediately placed in the insert. After being incubated at 37°C for 30 min, the inside of the insert was filled with Vas-cuLife at a higher medium level than the outside, resulting in a hydrostatic pressure difference and thus interstitial flow through fibrin hydrogels. The medium was refilled every day for over a 1-week duration.

The generation of microvascular network in Platform B was based on a previously established protocol (37). Briefly, dissociated HUVEC and NHLF were resuspended in a fibrinogen solution at 8 million cells/mL and 2 million cells/mL, respectively. With a pre-chilled pipette tip, 9 µL of cell-embedded fibrinogen solution was quickly mixed with 1 µL of 50 U*/*Ml thrombin in DPBS, resulting in the final concentration of 10 mg*/*mL fibrinogen and 5 U*/*mL thrombin, before being injected in the gel-loading chamber of the microfluidic chip. After sealing both gel-loading ports, the chip was then incubated at 37°C for 20 min. VascuLife was used in the co-culture and filled into each medium reservoir. On one medium channel, the inlet reservoir No. 1 was filled with 1500 µL (equivalent to 23 mmH_2_O) and the outlet reservoir No. 2 was filled with 500 µL (equivalent to 8 mmH_2_O). On the other channel, the inlet reservoir No. 3 was filled with 1000 µL (equivalent to 18 mmH_2_O) and the outlet reservoir No. 4 was filled with 200 µL (equivalent to 3 mmH_2_O). This medium-filling configuration ensures a hydrostatic flow along the medium channel (reservoir No. 1→ No. 2, and reservoir No. 3 →No. 4) as well as across the gel-loading chamber (reservoir No. 1&2 →No. 3&4). All medium reservoirs were replenished to the initial level every day over a 1-week duration, where the medium level of each side channel was reversed (reservoir No. 1&2↔ No. 3&4) each day to prevent HUVEC from migrating preferentially toward one side channel.

## 3. Results

### A. Baseline Platforms

Interstitial flow simulations were conducted under baseline pressure configurations for both platforms. Figure 2a shows that both platforms successfully generated IF velocities within the physiological range 0.1 µm*/*s to 11.0 µm*/*s (31) under baseline pressure conditions. Platform A exhibits uniform IF velocities throughout its porous domain, attributed to its simple geometry, constant cross-section and unidirectional flow. In contrast, Platform B shows spatial variability along the vertical midline. These results indicate that both platforms can achieve flow conditions suitable for vascular network formation, despite their differing geometries.

**Fig. 2.**
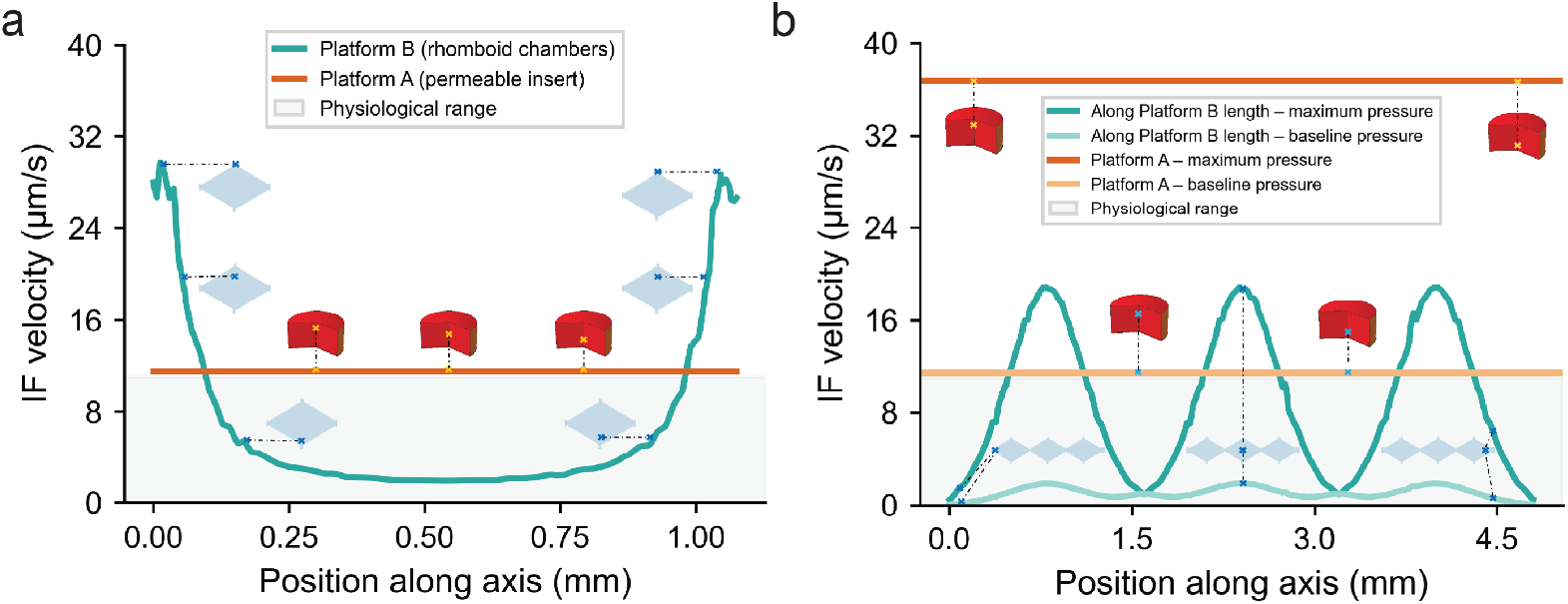
Interstitial flow (IF) velocity profiles extracted along vertical and horizontal midlines of the tissue chambers under baseline (16 mmH_2_ O) and increased (50 mmH_2_ O) pressure conditions. (a) Interstitial flow profile along vertical midline, baseline pressure. (b) Interstitial flow profile along horizontal midline at both baseline and increased pressures.

### B. Experimental Validation of Microvessel Formation

Over 7 days of culture, HUVECs, as our model endothelial cells, gradually moved and connected to one another to form a microvascular network across the entire hydrogel. Platform B exhibited an interconnected network of lumens or hollow channels inside. This network was perfusable with Dextran Rhodamine solution without any leakage into the hydrogel space, indicating the patency of the microvessel. Similarly, Platform A also showed the formation of microvascular network, though the overall diameter of the microvessels was smaller (Fig. 3).

**Fig. 3.**
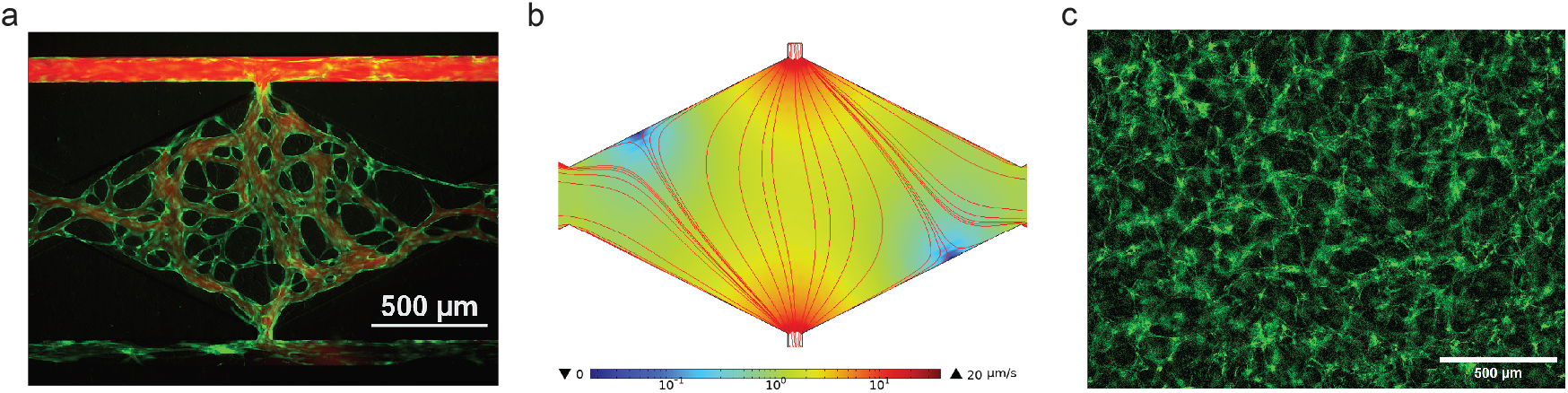
Experimental microvascular network formation in Platforms A and B, compared with CFD-predicted interstitial flow streamlines (Platform B only) (a) Microvascular network in Platform B perfused with Dextran Rhodamine after day 7 of culture. Flow direction is from top to bottom. (b) CFD-predicted interstitial flow streamlines (red) and magnitudes (colormap) for Platform B. (c) Bottom view of the microvascular network in Platform A after day 7 of culture, flow direction towards the reader.

Because interstitial flow (IF) is known to influence both the morphology and function of developing microvascular networks, we conducted CFD simulations using the same hydrostatic pressure configurations applied in the experiments. This allowed us to assess whether the observed vascular patterns aligned with predicted IF fields. The resulting interstitial flow (IF) field in Platform B showed a strong central convergence of streamlines (Fig. 3b). This spatial pattern corresponded closely to the experimentally observed vascular network occupying the same region (Fig. 3a). This correspondence between CFD-predicted flow and vascular morphology suggests that the IF field serves as a reliable indicator of whether a given geometry and perfusion condition will support successful microvessel formation. Therefore, CFD simulations can be used to evaluate the suitability of new designs or scaled-up platforms by examining whether their IF velocity falls within the range shown to be conducive to vascularization.

Figure 3c shows the resulting microvessel networks obtained in Platform A. Unlike Platform B, where imaging was performed in the same plane of the interstitial flow direction, images in Platform A were acquired perpendicular to the IF direction. This precludes a direct spatial comparison between predicted streamlines and vascular alignment.

### C. Controlling IF through pressure gradients

Scaling up platforms presents notable challenges, particularly the reduction in interstitial flow (IF) velocities when the pressure is kept constant. According to Darcy’s law, IF velocity is inversely proportional to the characteristic length, i.e., length measured along the main direction of the flow. As platforms are scaled up, this characteristic length might increase, leading to a natural decrease in IF velocities if the pressure difference driving the flow is kept constant. For a platform to remain viable during scaling, it must generate IF velocities above the physiological lower limit 0.1 µm*/*s, required for vascular network formation.

Figure 2b illustrates changes in IF velocity profile in both platforms under baseline pressure conditions (a hydrostatic pressure difference of 16 mmH_2_O) and the maximum pressure gradient we imposed as a design constraint (50 mmH_2_O). From this data, it is clear that pressure can be used to modulate IF magnitudes across the gel while preserving the underlying spatial flow pattern. This modulation is limited, however, by the mechanical integrity and compressibility of the gel and the susceptibility of endothelial and supporting cells to pressure stimuli.

### D. Scaling Up Platform A

The diameter and gel thickness of Platform A were scaled, first independently and then jointly, to assess scale-up effects on IF patterns. Note that modifications of the diameter and/or thickness of the gel can be feasibly effected through the use of PDMS rings of different sizes, of larger-diameter off-the-shelf permeable inserts, or by modifying the volume of gel added.

The full scaling results are summarized in Fig. 4. Figure 4a shows the results of scaling the diameter by factors of 2X, 5X, 10X, and 20X with respect to the baseline design, with corresponding volume increases by factors of 4X, 25X, 100X, and 400X, respectively. IF velocity remained constant regardless of the scaling factor when the applied pressure was unchanged. In contrast, Fig. 4b shows that increasing the gel thickness resulted in a significant decrease in IF velocity for each pressure condition. Figure 4d provides the pressure adjustments required to maintain constant IF velocity across various height scaling factors, offering practical guidance for preserving physiological flow conditions during scaling.

**Fig. 4.**
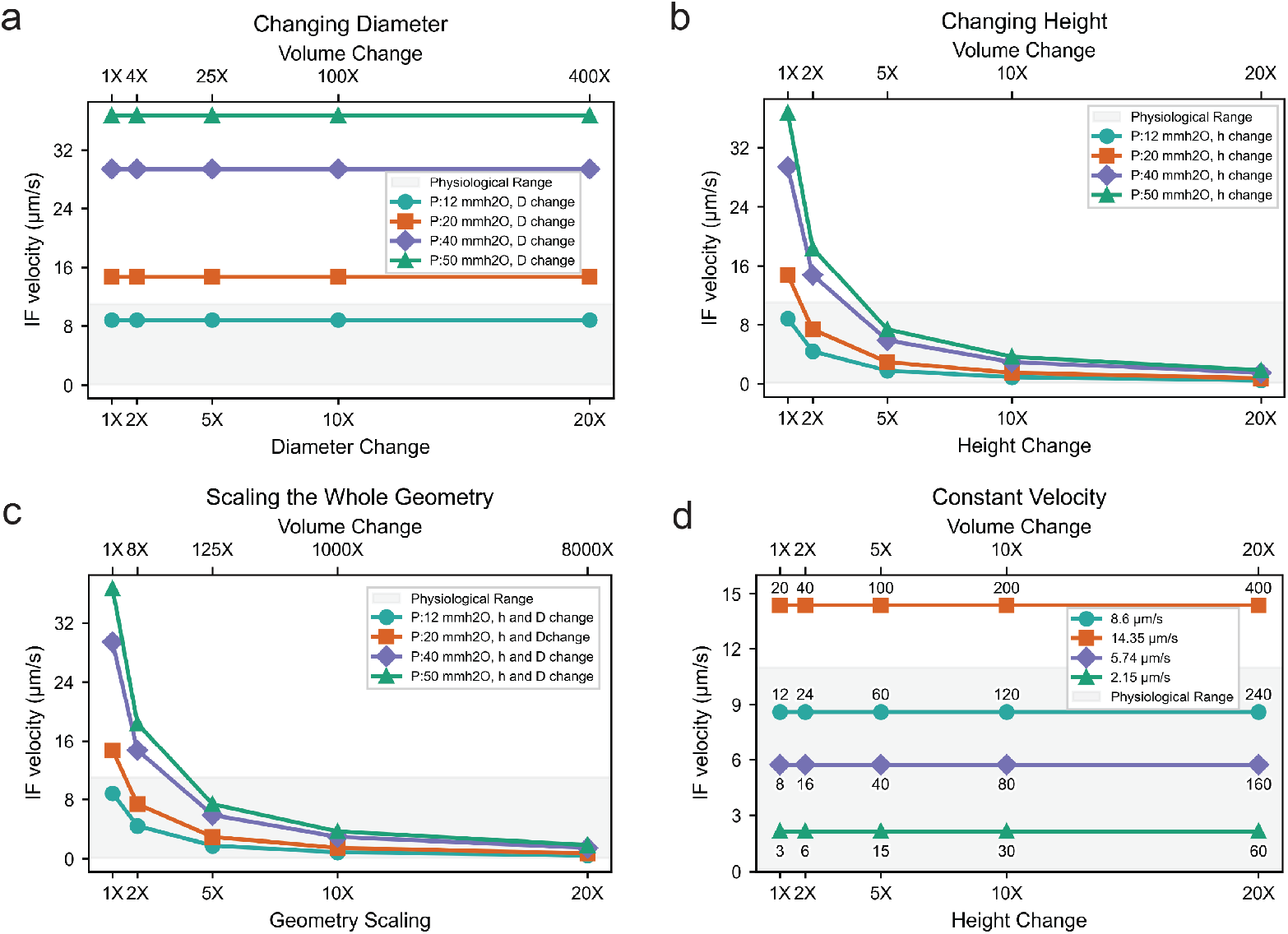
Scaling for Platform A (Permeable Insert): (a) Interstitial flow (IF) velocities for scaling the diameter, showing no change in velocity with increasing diameter under constant pressure. (b) IF velocity for scaling the height (gel thickness), showing significant decrease in velocity with increased height. (c) IF velocity for scaling the whole geometry (both diameter and height simultaneously), highlighting dominant effect of gel thickness on IF. (d) Required pressure adjustments (numbers above/below data markers) to maintain physiological IF velocity (colors) across various gel height scaling factors.

Additionally, the simultaneous scaling of both diameter and gel thickness was evaluated. As depicted in Fig. 4c, this scaling approach significantly reduces IF velocity across all pressure conditions, with gel height having the dominant influence. These findings provide guidance for dimension-specific scaling strategies, balancing IF control, increased yield, and experimental feasibility.

To validate these simulation results and demonstrate the feasibility of scaling up Platform A, microvessel formation was compared in permeable insert devices with three different gel volumes: a 25 µL baseline geometry and two scaled-up geometries of 200 µL (results not shown) and 800 µL. The increase in volume was achieved by enlarging the transwell diameter in Platform A. In all cases, the cell density and matrix formulation were kept constant at 9.5 M/mL HUVEC and 0.5 M/mL pericyte embedded in 10 mg*/*mL fibrinogen with 5 U*/*mL thrombin. GFP-expressing HUVECs formed interconnected microvessel networks in all conditions, as shown in Fig. 5.

**Fig. 5.**
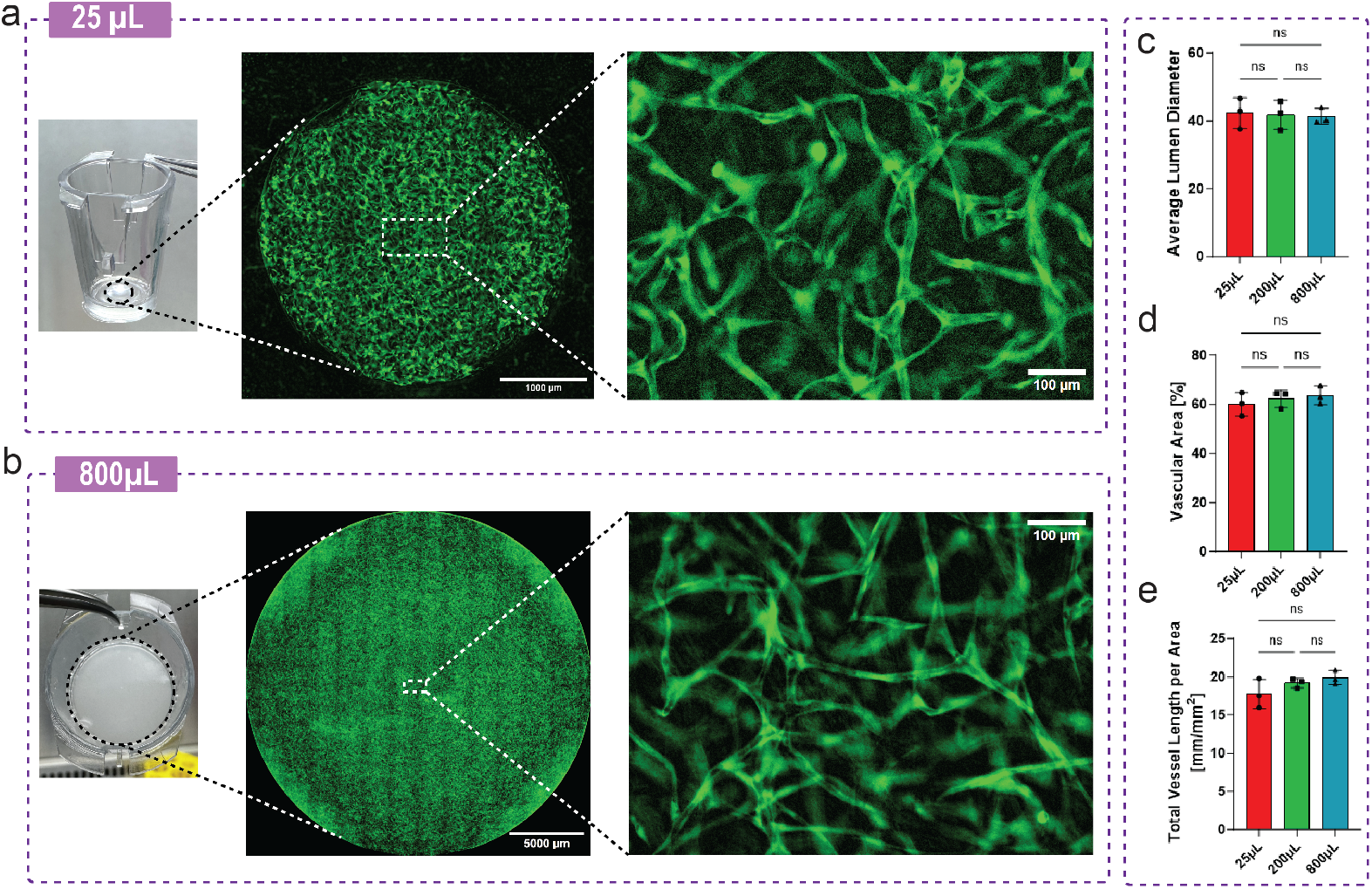
Microvessel formation and quantitative analysis in scaled-up permeable insert devices. (a–b) Left to right: photos of gel loading in 25 µL and 800 µL inserts, whole-gel fluorescence images (Day 5), and high-magnification views of GFP-HUVEC networks. Cell formulation: 9.5 M*/*mL HUVEC and 0.5 M*/*mL pericyte in 10 mg*/*mL fibrinogen with 5 U*/*mL thrombin. (c–e) Quantification of average lumen diameter, vascular area fraction, and total vessel length per area (mm*/*mm^2^) after 7 days of culture using 9 M*/*mL HUVEC and 1 M*/*mL pericyte with 10 U*/*mL thrombin. No significant differences observed between 25, 200, and 800 µL conditions (ns). Statistical tests: Shapiro-Wilk for normality; Brown-Forsythe and Welch ANOVA with Dunnett’s T3 post-hoc test (*n* = 3/group).

To assess the effect of scaling on network morphology, vessel characteristics were quantified at Day 7. Statistical analysis was performed using the Shapiro-Wilk test for normality, followed by Brown-Forsythe and Welch ANOVA with Dunnett’s T3 multiple comparisons test to account for unequal variances. Across all groups (n = 3 per group), no statistically significant differences were found in average vessel diameter, vascular area, or vessel length density. This indicates that interstitial flow, which remained constant across bioreactor scales, effectively modulates microvessel network topology in this setting.

### E. Scaling Up Platform B

Unlike Platform A (Permeable Insert), Platform B (rhomboid) does not exhibit uniform interstitial flow (IF) velocity throughout its domain. Due to spatially varying nature of IF in this platform, box-and-whisker plots were used to represent the IF distribution across different scaling strategies. Figure 6 presents an array of results for independent scaling of different dimensions of Platform B (figure rows), namely length and width (distance between inlet and outlet of the chambers), under varying pressure conditions (figure columns). It can be observed that as the scaling factor increases, the overall range and inter-quartile range (IQR) of the IF distributions decreases with respect to the baseline geometry (1X). At larger scaling factors, the system creates a narrower range of IF velocities compared to the baseline geometry, particularly under lower pressure conditions (e.g., 12 mmH_2_O). It can also be observed that across scales, a portion of the gel volume is exposed to IF outside the target range (shaded area in the plots), specially at larger pressures.

**Fig. 6.**
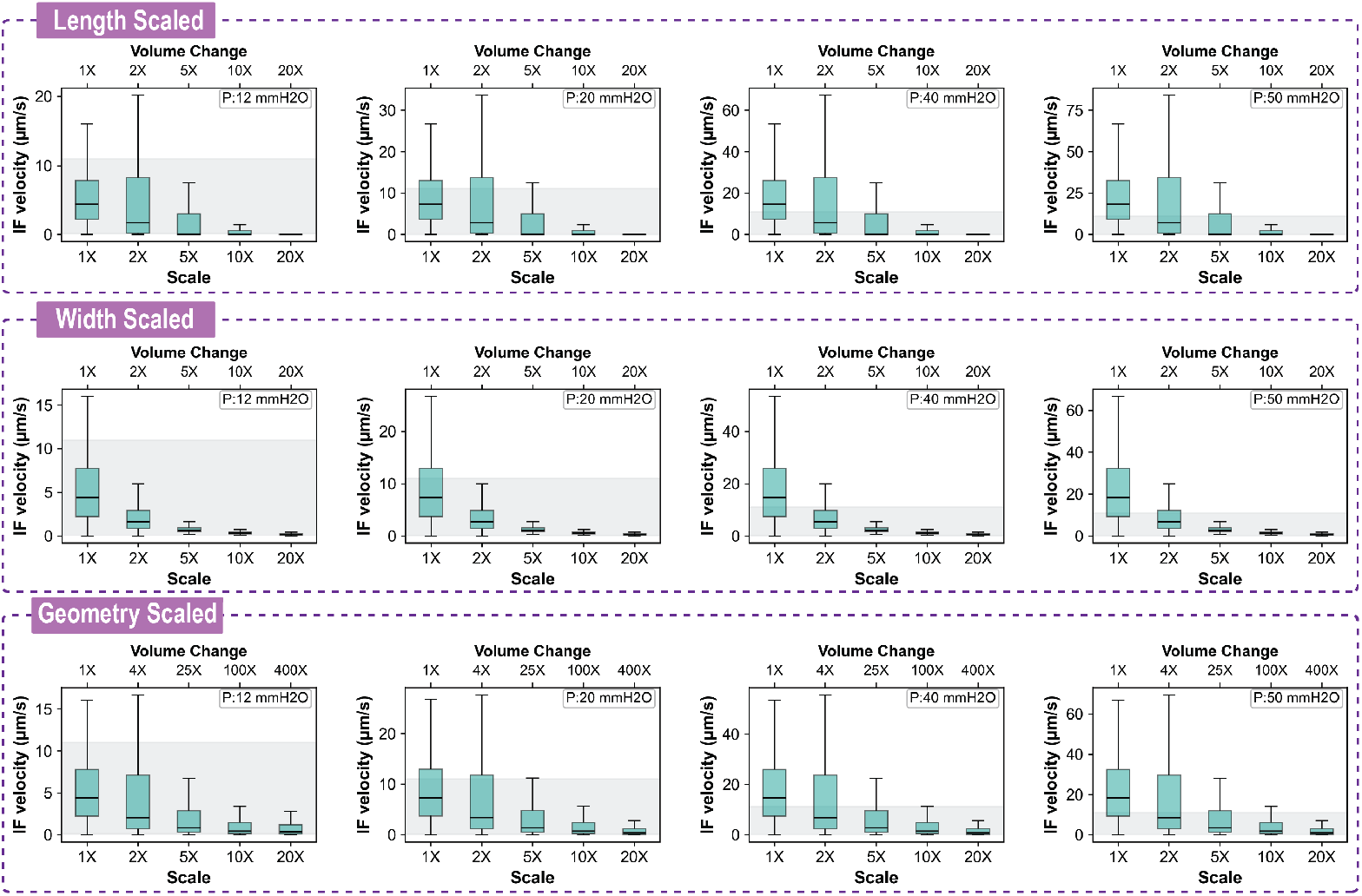
Interstitial flow velocity distribution for scaling different dimensions of Platform B (rows) under varying pressure conditions (columns). Shaded area in all figures indicates the physiological interstitial flow range, which has been shown to lead to microvessel network formation in vitro. Baseline geometry corresponds to 1X.

The third row of plots in Fig. 6 presents the results of scaling the entire rhomboid geometry with a constant aspect ratio. Compared to scaling individual dimensions, scaling the entire geometry provides significant increases in chamber volume while still replicating the IF distribution of the baseline geometry. The results indicate that scaling the entire geometry up to 5X maintains interstitial flow (IF) velocities within the physiological range across the domain. Beyond this scale factor, the velocity distribution narrows compared to the baseline geometry. These findings suggest that whole-geometry scaling is effective up to moderate scale factors, but maintaining the IF distribution at larger scales requires adjustments in the applied pressure gradient.

### F. Geometry Comparison Under Controlled Volume and IF Conditions

Platform A (permeable insert) and Platform B (rhomboid chamber) were compared to evaluate how their geometries influence IF velocity distribution under matched gel volume and average IF velocity conditions. IF velocities for the baseline design were extracted, and Platform A was scaled to match the total gel volume of the baseline Platform B. Pressure adjustments were applied to achieve the same average IF velocity in both platforms. Figure 7 illustrates the IF velocity distributions for the two platforms. Platform A generates a uniform IF velocity field across the gel, as indicated by the horizontal red line in the Fig. 7a (also the vertical red line in Fig. 7b). In contrast, Platform B exhibits a wide range of IF velocities across different regions of the gel, as shown by the box-and-whisker and histogram plots. The variability in Platform B highlights its ability to create localized flow profiles, while Platform A maintains consistent flow throughout its domain. Note that these IF distributions can be correlated to the diameter or length of the resulting microvessel networks. Figure 7c and Fig. 7d show the vessel characteristics that were observed for different interstitial flow values by Zhang et al. (13).

**Fig. 7.**
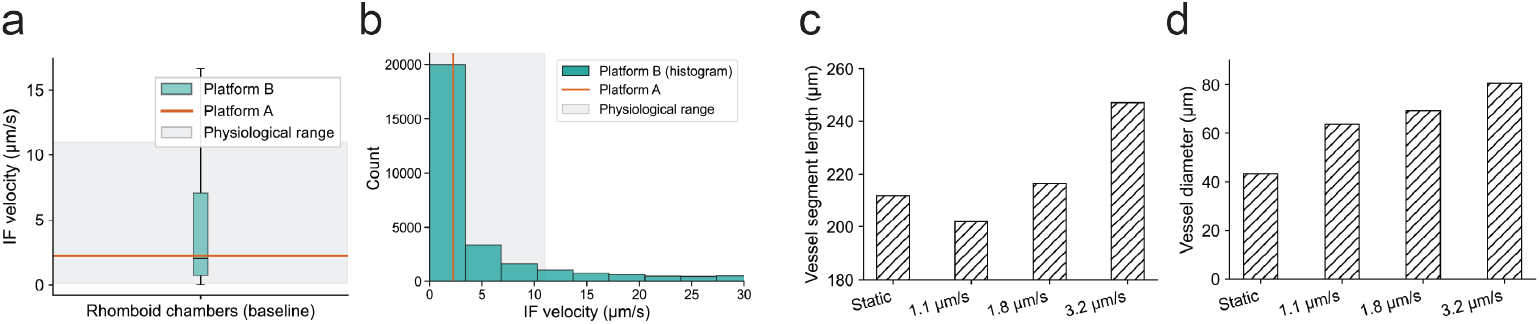
Interstitial flow (IF) velocity distributions for Platform A and Platform B under matched gel volume (1.1 mm^3^) and average IF velocity (2.2 µm*/*s) conditions. The shaded region represents the physiological IF range 0.1 µm*/*s to 11.0 µm*/*s. (a) Boxplot of IF velocities, (b) Histogram of IF velocities, (c) and (d) Microvessel lengths and diameters obtained at different IF velocities by Zhang et al. (13).

## 4. Discussion

This study explored the scalability of Platform A (permeable insert) and Platform B (rhomboid chambers), focusing on interstitial flow (IF) velocities across various scale-up factors and pressure conditions. While both platforms demonstrated scalability potential, Platform A exhibits greater ease of control and spatial uniformity in IF velocity regardless of scaling, with scaling options that do not require larger pressure gradients to sustain a target IF. In contrast, Platform B exhibits a wide range of IF, and requires pressure adjustments to sustain a target IF under most scaling scenarios. Experimental validation results suggest that IF is the main enabling factor for generation of microvessel networks, at least within the target IF range used in this work. While the significance of IF gradients on network formation was not characterized herein, it was shown by Staples et al. (38) that endothelial cells predominantly polarize against the direction of flow at high shear stresses (0.1 Pa), while up to 55% of endothelial cells polarize in the direction of the flow, or even orthogonally, at lower shear stress levels (0.08 Pa). Importantly, shear stress has also been shown to modulate intususceptive angiogenesis (38) and angiogenic sprouting (39). Our results (e.g., Fig. 3a and Fig. 3b) suggest that the presence of IF gradients modulates the diameter and connectivity of the resulting network. Note that microvessels generated in Platform A, where no significant IF gradients occur, show remarkably similar diameters, in contrast with those obtained in Platform B.

### A. Comparative Insights on Scaling

The uniform IF velocity profile of Platform A aligns with previous findings (12), which confirms its suitability for scaling scenarios where consistent flow conditions are essential. For Platform B, scaling revealed significant variations in IF velocity across different areas of the geometry, including at relatively small scale factors. This was confirmed by Dextran perfusion experiments (e.g., Fig. S1), showing uneven gel saturation, consistent with spatially varying IF velocity. The spatial gradients of IF may more closely mimic physiological conditions leading to formation of vascular networks. Indeed, several studies reporting successful microvessel formation employed platforms that inherently exhibit spatial IF gradients due to their geometries or perfusion strategies (13, 18, 31, 37), leading to non-uniform vessel diameter distributions.

Recent evidence indicates that IF gradients in interstitial flow (in both magnitude and direction) can modulate vascular network morphology. Controlled 3D models have demonstrated that mild IF levels (0.1–5 µm/s) promote greater vessel area, branch length, and average diameter compared with static cultures (40), with flow gradients on the order of 0.01–0.1 µm/s per 100 µm correlating with directional bias in sprouting (18). Convective transport also reshapes morphogen distributions, as low physiological IF (0.5–5 µm/s) can dissipate VEGF gradients, redirecting sprouting against the flow vector and reducing local branching density (31). Collectively, these findings suggest that moderate IF gradients (roughly 0.01 – 0.05 µm/s per 100 µm spatial scale) foster complex, branched vascular topologies, whereas steeper gradients bias toward enlarged, mature vessel structures. It is unclear, however, if IF gradients affect network morphology directly or, instead, indirectly through their effect on morphogen (e.g. VEGF) gradients. Regardless of the underlying mechanism, the broader IF spectrum observed in Platform B may produce networks with greater variability in diameter and topology, while Platform A’s uniform IF field is suitable to yield more homogeneous architectures, as was indeed shown in our experimental results. Purposeful engineering of gentle IF gradients—through graded permeability or tapered geometries—could therefore serve as a design lever for tailoring microvascular morphology in scalable bioreactors.

In addition, results from Platform B suggest the existence of an upper boundary for effective scaling, beyond which achieving both physiological IF velocities and a flow distribution conducive to microvessel formation becomes more challenging. Tailored scaling strategies aligned with specific experimental objectives are therefore essential to optimize scalability. For example, scaling the width of Platform B proved less effective as a standalone strategy, requiring higher pressure adjustments to achieve target velocities. Scaling the entire geometry, however, demonstrated a better balance between volume increases and maintaining IF velocity, although a lower inter-quartile range of the IF distribution was observed at higher scaling factors, potentially impacting the diameter, length or connectivity distributions of the resulting microvessel networks, as was observed by Soon et al. (37).

Experimental validation in the scaled-up versions of Platform A confirmed the formation of perfusable microvascular networks under hydrostatic pressure-driven flow. Additionally, quantitative assessment of vessels formed within Platform A across scaled-up diameters showed consistent vessel diameter distributions, vascular area, and vessel length per area across different gel volumes, despite a more than 30-fold increase in scale. This consistency, achieved while maintaining constant interstitial flow conditions, shows the pivotal role of IF in regulating microvascular morphology. Successful vessel formation under IF, along with quantitative morphological assessment, has also been demonstrated in Platform B (37).

### B. CFD-based Scale-Up

This study demonstrated the feasibility of scaling up Platform A (Permeable Insert) and Platform B (rhomboid) while maintaining IF within a target range using numerical simulation approaches. CFD simulations provided a detailed understanding of spatially varying IF profiles in the examined platforms, and elucidated how pressure adjustments can maintain functional flow conditions during scaling. Combining CFD and in vitro results, this works provides further evidence of the role of IF in enabling the formation of microvessel networks and controlling its characteristics. Our scale-up results on Platform A, where we imposed the same IF on multiple scales of the same geometry, showed remarkable uniformity. In contrast, the spatially varying IF profile obtained in Platform B, resulting in a distribution of microvessel diameters and lengths, a finding that has been reported in multiple previous works (13, 17, 18). The observed qualitative agreement between predicted flow patterns and experimentally formed microvessel networks further reinforces the utility of CFD as a predictive tool for design optimization.

Importantly, with further evidence that IF modulates microvessel network characteristics, IF distributions predicted with experimentally validated CFD models can be used to obtain optimal bioreactor designs to meet any IF target range, and thus any target distribution of vessel diameters or lengths.

### C. Simulation-based Design Optimization

To illustrate this point, we formulated a simulation-based design optimization problem to modify the design of Platform A. Specifically, we modified the insert geometry to have several cross-sections, similar to a set of vertically stacked cylinders with different heights and radii, joined by short transition sections shaped like inverted truncated cones, as shown in Fig. 8a. We defined as design parameters a) the height *Hi* and radius *ri* of each vertical section, and the height *hi* of the transition sections. We conducted a dimensional optimization study in COMSOL, with the same simulation settings described in the Methods section, and with an inlet pressure of 68.6 Pa (7.0 mmH_2_O) applied on the top of the insert, with the goal of achieving a target distribution of interstitial flows inside the gel volume.

**Fig. 8.**
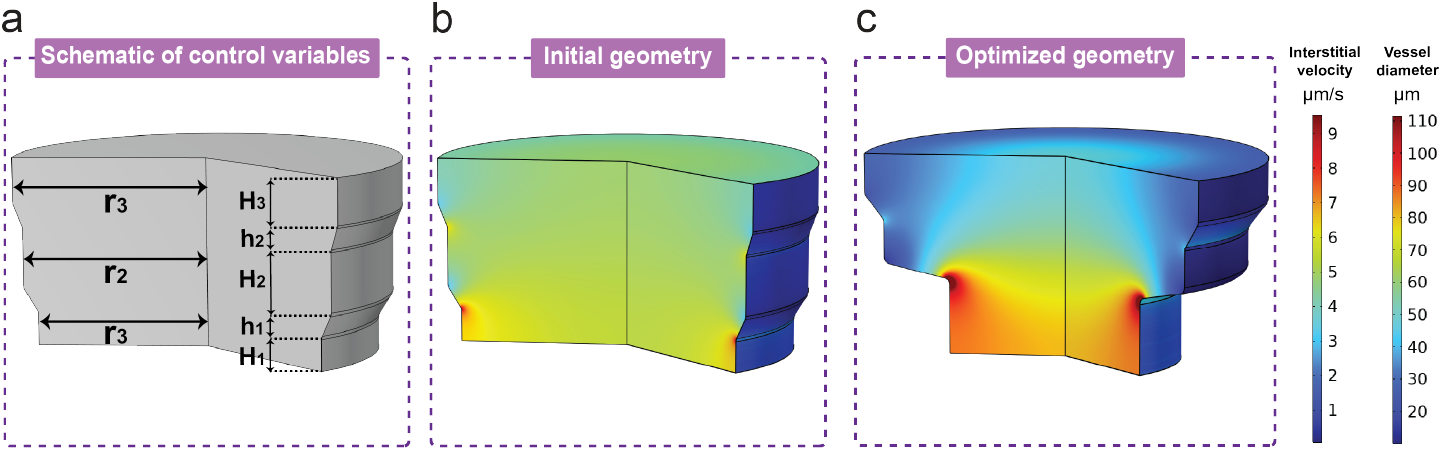
Optimization of modified Platform A for targeted IF distribution. (a) Design variables, (b) IF velocity field in the baseline design, corresponding to *x*_0_ = (0.3500, 0.6500, 0.5000, 1.8000, 1.9500, 2.0500, 0.2500, 0.2500) × 10^−3^ m, (c) IF velocity field in the optimized design, corresponding to *x*_*opt*_ = (0.7468, 0.4318, 0.4139, 1.1519, 1.8052, 1.9938, 0.1545, 0.2531) × 10^−3^ m. Maximum and minimum values in the color scales shown on the right of the figure, corresponding to IF velocity and the expected microvessel diameter (Fig. 7d), have been clipped to facilitate the simultaneous depiction of IF and expected vessel diameter in the same figure.

Let **x** = [*H*_1_, …, *H_N_*, *r*_1_, …, *r_N_*, *h*_1_, …, *h*_*N* −1_] be the vector of design parameters. Let *I* = *I*_1_, …, *I_N_* be a partition of the target IF range [0.1, 15]µm*/*s into *N* intervals,

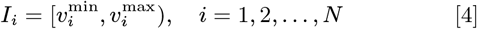

where 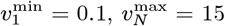, and 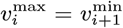 for all *i*. For each IF interval *Ii*, we compute the fraction of the gel volume with IF in the corresponding range, *f_i_*(**x**), and we denote the target fraction for that interval as 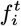. Then, the optimization problem is formulated as

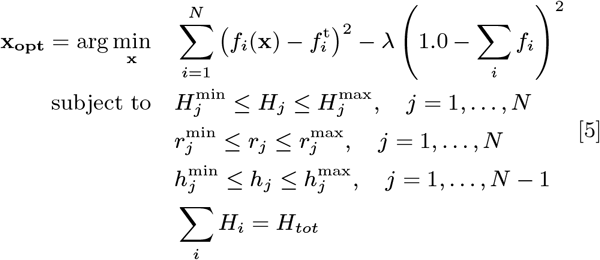

where *H_tot_* is the total height of the permeable insert. Note that the objective function is the sum of squared deviations from the targets and includes an additional term that seeks to minimize the fraction of the gel with IF velocities outside the target ranges.

For illustration purposes, let us define three target IF ranges, namely 0.1 µm*/*s to 1.0 µm*/*s, 2.0 µm*/*s to 3.0 µm*/*s, and 5.0 µm*/*s to 10.0 µm*/*s, corresponding to three vertical sections for the permeable insert. Table 1 shows the design parameter ranges and the rationale for our choices.

**Table 1.** Design space for optimization of modified Platform A. Three gel zones are defined, separated by two transition zones, to meet three different target IF ranges with a constraint on the maximum height of the system. Design ranges for radii and heights are defined around the baseline design, enforce a constant system height, and non-vanishing top zone and transition zones. The section and transition heights satisfy *H*_1_ + *h*_1_ + *H*_2_ + *h*_2_ + *H*_3_ = *H*_tot_ = 2.00 mm.

| Design Variable | Baseline | Step | Min | Max | Rationale |
| --- | --- | --- | --- | --- | --- |
| Bottom radius $r_1$ | 1.80 mm | 0.002 | 1.00 mm | 2.15 mm | Maximum radius is baseline 2 mm + buffer, minimum radius keeps sizes reasonable. |
| Middle radius $r_2$ | 1.95 mm | | | | |
| Top radius $r_3$ | 2.05 mm | | | | |
| Bottom section height $H_1$ | 0.35 mm | 0.001 | 0.30 mm | $\leq 1.40$ mm* | Total height is baseline 2 mm, Maximum and minimum set to enforce 3 zones with 2 smaller, non-vanishing transitions. |
| Middle section height $H_2$ | 0.65 mm | | | | |
| Top section height $H_3$ | 0.50 mm | | | | |
| Bottom transition height $h_1$ | 0.25 mm | 0.001 | 0.15 mm | 0.30 mm | |
| Middle transition height $h_2$ | 0.25 mm | | | | |

Figure 8 shows the results of this optimization study. Specifically, Fig. 8b and Fig. 8c depict the geometry and IF flow field of the initial and optimized designs, respectively. As expected, successive reductions of cross-sectional area cause the flow to accelerate, resulting in a spatially varying IF distribution. In addition, the changes in cross-section modify the direction of the flow, introducing radial velocity components, as shown by the flow streamlines and velocity vectors depicted in the supplementary material, Fig. S2.

Studies by Abe et al. (17) and Kim et al. (18) have shown that flow gradients and directionality can bias the orientation and extension of angiogenic sprouts, with angiogenesis often occurring opposite to the direction of flow. Therefore, CFD-based design of the bioreactor geometry, which allows us to control the spatial distribution of IF magnitude, direction and gradients, can enable the design and scale-up of bioreactors for microvessel formation with targeted topology and morphology. For instance, given a mapping correlating IF velocities with the geometrical and topological characteristics of the resulting microvessel networks, the optimization problem of Eq. 5 could be recast in terms of target distributions for vessel diameter, vessel length, total vessel area coverage and/or average number of branching points (17). In this work, we used experimental results by Zhang et al. (13) that characterize the average vessel segment length and diameter as a function of the IF used during the culture period (Fig. 2 in the reference). Based on this data, we fitted a linear regression to map the IF fields depicted in Fig. 8b and Fig. 8c, to fields of predicted vessel diameters, please refer to the rightmost colorbar in Fig. 8.

This study illustrates the value of CFD models for bioreactor design. As the effects of the mechanical and biochemical environment on microvessel formation and morphology are elucidated, CFD models, potentially augmented with the inclusion of mathematical models for cell signalling networks (41), could enable the manufacture of microvessels at scale, with accurate, robust control of the resulting microvessel network characteristics.

### D. Limitations and Future Studies

In this study, we assumed uniform, average permeability and porosity throughout the gel. Thus, our CFD simulations may not capture the spatial variability that may be present in manufactured gels, which would impact IF velocity distributions. Future research may incorporate detailed models derived from high-resolution image analysis, such as segmented z-stacks (e.g., Zhao et al. (42)).

These models would enable more accurate predictions of local IF velocity. Specifically, predicting pore-scale velocity and shear stress would better describe the mechanical environment directly experienced by the cells. Combined with multi-scale modelling (41), such approaches could predict spatio-temporal variations in IF, permeability, porosity and microvessel formation.

Compressibility of the fibrin gel under pressure was not considered. This could potentially affect porosity and permeability, particularly under high pressure. However, our initial in silico tests indicate that accounting for gel compressibility did not impact IF predictions. Specifically, we observed *<* 0.1% change in predicted IF, at an inlet pressure of 127.5 Pa (13.0 mmH_2_O), for a gel with elastic modulus of 0.9 kPa and Poisson’s ratio of 0.49. However, fibrin gel stiffness is known to modulate directional angiogenic bias (31), an effect not considered in our hydrodynamically focused study. Similarly, we did not consider potential effects of pressure stimuli on endothelial cell function, signalling and gene expression (43). Finally, this study focuses on IF velocity as the key descriptor, leaving out shear stress (39, 40) or matrix evolution. Future work should expand this to consider shear stress and dynamic remodelling effects, such as matrix compaction and permeability evolution, which alter local flow and nutrient distribution and thus impact vessel network characteristics.

## 5. Conclusions

This study provides a comprehensive assessment of the scalability of two published platforms for production of microvessel networks, intended for co-culture and pre-vascularization of engineered tissues. CFD simulations of the baseline geometries, combined with in vitro perfusion experiments on fibrin/thrombin gels seeded with HUVECs, along with fibroblasts and pericytes, were used to characterize the flow conditions, quantified in terms of interstitial flow velocity, under which microvessels form, in agreement with previous literature. Subsequently, CFD simulations were used to study the interstitial flow fields for scaled-up versions of the two platforms, where scaling factors were applied independently to each dimension, as well as to both dimensions simultaneously. Results demonstrated the feasibility of scaling up these platforms while maintaining IF conditions within the target ranges consistent with physiological values. Experimental validation of Platform A scale-up confirmed that microvessel formation can be maintained across a more than 30-fold scale increase when IF conditions are preserved. Prior studies on Platform B also support the formation of functional microvessels under pressure-driven IF, underscoring the broader relevance of the findings. This correspondence, alongside flow-field agreement with CFD predictions, reinforces the practical applicability of the modeling framework for design and optimization of bioreactors for microvessel production.

Scale-up results for the two platforms indicate that Platform A exhibits uniform flow profiles, with direct control of the pressure gradient driving the flow, with microvessel networks with vessel diameters that are remarkably consistent across scales for any given IF. For instance, scaling Platform A diameter by 10X results in a 100-fold increase in volume (i.e., yield) while sustaining the same IF conditions as the baseline without requiring an increase in the pressure gradient. In contrast, Platform B exhibits a spatially varying IF profile that more closely mimics the complex IF environments of native tissues, resulting in vessel networks with a wide variety of vessel characteristics such as diameter, length and connectivity.

By combining computational modelling with experimental validation, this study provides a framework for evaluating and scaling perfusion bioreactors systems based on interstitial flow dynamics.

## 7. Data availability

The data underlying this article will be made available after acceptance for publication in an archival repository (Zenodo).

