## Supplementary Material for "Computational fluid dynamics enables predictable scale-up of perfusion bioreactors for microvessel production"

February 11, 2026

### 1 Porosity estimation

To characterize the gel used for the experiments, the gel chamber of the Platform B was filled with a fibrin hydrogel containing 10 mg/mL fibrinogen (F8630, Sigma Aldrich) and 5 U/mL thrombin (T4648, Sigma Aldrich). The platform was left at room temperature for 15 min before being further incubated at 37 °C in a $CO_2$ humidified incubator to polymerize the hydrogel. Then 200 µL of prewarmed DPBS(-/-) was added to each medium side-channel to completely wet the gel-port interfaces and remove gas bubbles. Subsequently, 1 mL of 70kDA neutral Dextran Rhodamine B (ThermoFisher, D1841) solution in DPBS(-/-) at 100 µg/mL final concentration was added to one of the media inlets, resulting in 23 $mmH_2O$ hydrostatic pressure difference to the other three inlets/outlets. The resulting hydrostatic pressure led to the development of a perfusion flow and thus created a gradient of Dextran concentration across the fibrin hydrogel over time. Immediately after adding the Dextran solution, Z-stack images were taken every 2.5 min with a widefield fluorescent microscope (Axio Observer Z1, Carl Zeiss). Each image was 2048 × 2048 pixels across 2.66 mm × 2.66 mm. Figure S1a shows the Dextran-perfused chamber at times 0 min and 35 min, the smaller panels depicting the perfusion dynamics every 5 min.

Figure S1b shows a sample image of the gel chamber 35 min after perfusion started, with a different colormap. The microchannel on the left (yellow-coloured region) is the inlet to the chambers, and thus this pixel image intensity level corresponds to regions fully occupied by the Dextran solution. Similarly, on the right, there is a barely visible outlet microchannel that has not been perfused with Dextran solution. We use an area inside the rhomboid chamber but close to the outlet as the reference value for zero Dextran occupancy. Figure S1c depicts an enlargement of an area of the gel chamber located 10 µm from the inlet. This image clearly shows an array of pixel intensities, corresponding to different levels of Dextran occupancy of the gel volume covered by that image pixel. Since at this time point (35 min after perfusion) Dextran has clearly saturated the inlet region of the chamber, the variations in image intensity seen in the panel correspond to different proportions of Dextran and gel fibers present in the gel volume captured by the pixel, and thus form the basis for our estimation of porosity.

Porosity values were estimated by linearly interpolating between the image intensity values corresponding to full Dextran occupancy (inlet microchannel, no gel) and zero Dextran occupancy (fibrin gel near the chamber outlet, no Dextran). The implied assumptions are that a) fibrin gel fibers attenuate light proportionally to their occupancy

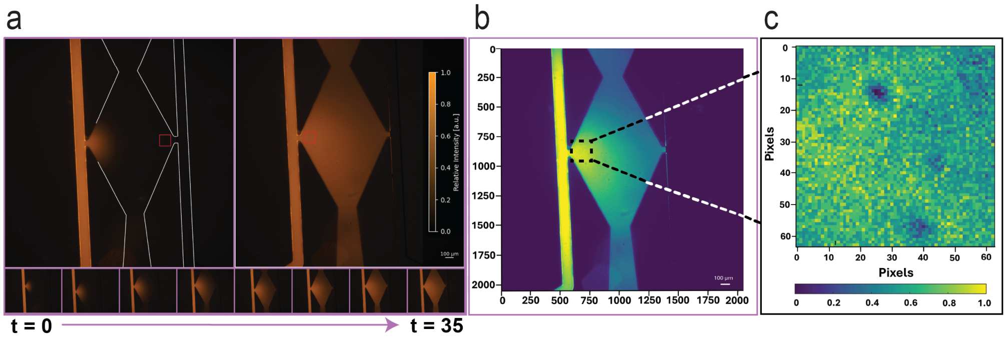


Figure S1: Dextran perfusion of endothelial-seeded gel in Platform B. (a) Time sequence of perfusion dynamics, images taken every 5 min until 35 min, the initial ($t = 0$ min) and final ($t = 35$ min) images are shown enlarged, (b) Top view of a chamber 35 min after initiation of perfusion. (c) Enlargement of an area near the inlet channel. Each pixel in the image corresponds to an area approximately 1.29 µm × 1.29 µm.

within the voxel corresponding to a given pixel, b) voxels corresponding to pixels are sufficiently large relative to fibrin fibers, and c) the camera sensor's response to the integral of light intensity over a pixel is linear. With these assumptions, the porosity is estimated as the average of the normalized pixel-level image intensity values over the area near the chamber inlet indicated in Fig. S1b, that is

$$\epsilon = \frac{1}{N} \sum_{i,j \in \mathcal{A}} \frac{I_{i,j} - I_{\min}}{I_{\max} - I_{\min}}, \tag{1}$$

where $I_{i,j}$ is the image intensity of a pixel with image coordinates $(i, j)$, $N$ is the total number of pixels in the averaging area $\mathcal{A}$, $I_{\max}$ is the intensity value corresponding to full Dextran occupancy ($\epsilon = 1.0$), and $I_{\min}$ is the intensity corresponding to zero Dextran occupancy ($\epsilon = 0.0$). Based on our experimental data (e.g., Fig. S1), the estimated porosity of the fibrin gel is 0.91.

### 2 CFD-based Platform Optimization

Figure S2 shows the velocity components, velocity vectors and streamlines of the baseline and optimized Platform A. Note that the changes in cross-sectional area cause flow acceleration and change the direction of the flow near the wall, generating radial velocity components. These changes in the direction and magnitude of the IF velocity have been correlated by previous work to the morphology (length, diameter), connectivity (number branching points) and preferential growth direction of the microvessel networks.

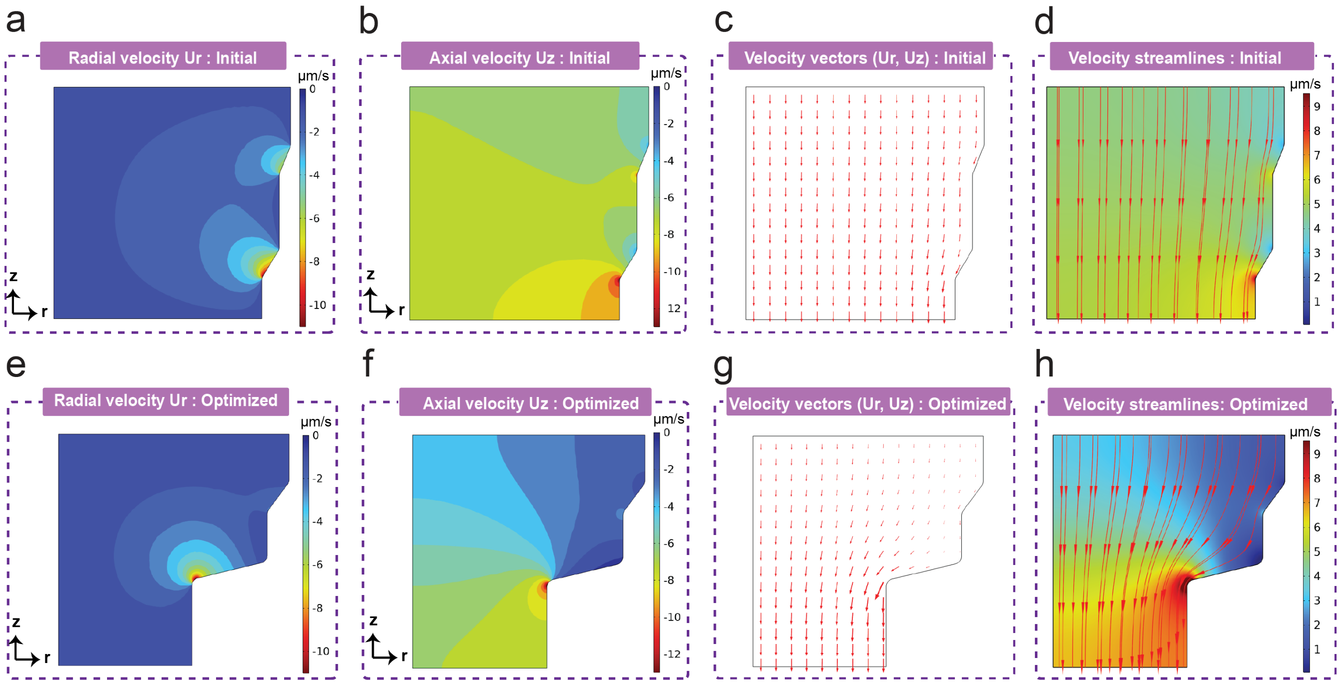


Figure S2: Interstitial flow field characteristics in optimized Platform A. Top and bottom rows show results for the baseline and optimized platforms, respectively. a) & e) radial velocity, b) & f) axial velocity, c) & g) velocity vectors, d) & h) streamlines (lines and arrows) and velocity magnitude (color flooding).
